# Repetitive stimulation modifies network characteristics of neural organoid circuits

**DOI:** 10.1101/2025.01.16.633310

**Authors:** Siu Yu A. Chow, Huaruo Hu, Tomoya Duenki, Takuya Asakura, Sota Sugimura, Yoshiho Ikeuchi

## Abstract

Neural organoids form complex networks but lack external stimuli and hierarchical structures crucial for refining functional microcircuits. In this study, we modeled the hierarchical and modular network organization by connecting multiple organoids and tested if the connection enhances the external stimuli-induced network refinement. We cultured networks of one, two, or three organoids on high-density microelectrode arrays, applied repetitive stimulation at two input locations from the microelectrodes, and monitored emergence of output signals that can decode the stimulus locations with machine learning algorithms. After two weeks of daily stimulation, networks of three organoids showed significantly higher stimulus decoding capability compared to the simpler one- or two-organoid networks. Long-term stimulation induced pronounced changes in the three-organoid network’s response patterns, spontaneous activity, and inter- and intra-organoid functional connectivity. These findings underscore the importance of hierarchical network organization, e.g. creating distinct subnetworks with specialized roles, for stimuli-induced formation of circuits with robust input-output functionality.

## Introduction

The brain is composed of diverse types of neurons and other cells that are intricately arranged to receive and interpret various signals from the world, enabling execution of complex functions such as thought, perception, and movement. Recent development in stem cell research and developmental biology rapidly advanced *in vitro* tissue modeling of the brain by allowing us to generate neural tissues—known as organoids—that recapitulate many of the key cellular and structural features of the brain^1,2^. While these organoids exhibit some fundamental brain-like features, their capacity to process external signals is far less than the brain^3–5^. This limitation stems from their relatively simple and disorganized neural circuits, which fall short of the brain’s sophisticated connectivity. However, this limitation offers a valuable opportunity to build up on current neural organoids to explore and understand mechanisms and requirements for functionality of neural circuits in vitro.

Broadly, there are three major steps to circuit formation in the brain^6,7^. First, cellular diversity and microscopic organization of brain regions are established. Local circuitries can be generated within the regional tissues. Second, functional regions are interconnected by tracts of axon. This allows them to be connected hierarchically, which determines major pathways for the signals and the stream of information within the brain. Finally, the neural circuits are further shaped and refined by exposure to complex activity^8–10^. Waves of neural activity, triggered spontaneously or externally, propagate through the neural circuits and induce ensemble of long-term plasticity programs within the cells which collectively shape functional circuits^9,11^. Exposure to the spatiotemporally diverse activity generate circuits that can distinguish different input signals and discriminating them, serving as the basis for processing information. Stimulus triggers response that encodes the information of the input signal, thus the spatiotemporal sequence of output signal from well-trained neuronal networks should be characteristic enough to be decodable^12^. Therefore, ability of biological neural network to interpret signal could be assessed by evaluating the “decodability” of its output signals (e.g. how reliably different inputs can be discriminated in the output signals that are intrinsically noisy).

Diverse cell types and three-dimensional structures of neural organoids form intrinsically complex neural networks which mimics the initial steps of circuit formation in the brain^13–16^. However, neural organoids do not receive external stimuli which is critical for organization and refinement for constructing functional biological neural networks. They also lack the hierarchical macroscopic circuit structures which are the key elements of the brain functionality^17,18^. In remains unclear, therefore, neural organoids are adequately equipped with cellular networks that can efficiently acquire functional circuits responsive for extrinsic signals. We hypothesized that connecting multiple organoids enhance their ability to establish long-lasting circuit alterations that enable the tissues to distinguish input signals. Multiple methods have been proposed to generate multi-organoid or multi-region circuit tissues. By fusing multiple organoids, multi-region interactions have been modeled^19–21^. We previously reported that two organoids connected by axons generated complex activity and displayed short-term plasticity^20^. When multiple organoids are interconnected, each organoid can serve as a modular unit—a subnetwork within a larger integrated network. By assigning specialized roles to the modular subnetworks, the network could gain hierarchical organization during functional assessment in vitro. However, it is still unclear whether such circuit structures can enhance their stimulation-induced refinement.

In this research, we investigated the effects of long-term repetitive stimulation on sparsely connected networks of cerebral organoids using stimulus decoding task as a functional assay. We cultured one, two, or three organoids on a high-density microelectrode array. The organoids on a microelectrode array extend axons, and if any, connect with other organoids, forming a sparsely but functionally connected neural organoid circuits. We stimulated the organoids at two distinct locations as input signals and monitored the response activity at another area of the organoids as output signals to evaluate if the source input can be discriminated blindly with the output signal (stimulus decoding task). We initially selected the input/output electrode locations so that a Support Vector Machine (SVM) algorithm cannot blindly distinguish the two source inputs from the output signal patterns. To evaluate long-term effect of input signals, we stimulated the organoids more than two weeks every day. Strikingly, the network of three organoids exhibited significantly greater SVM classification score after the long-term repetitive stimulation, suggesting the source signal can be discriminated with the output signal, while we did not observe the increase of the score for the neural network of one or two organoids. Similarly, CNN classification score also increased significantly with the network of three organoids. We found that the repetitive stimulation modified the response to stimulation of the network of three organoids. Spontaneous activity was also modified following the long-term stimulation in the network of three organoids. We observed that both inter- and intra-organoid functional network connectivity was altered by the repetitive stimulation in the network of three organoids. The results demonstrate the importance of circuit structure and connectivity in functionality of organoids and demonstrated the potential of connected organoids in future applications.

## Results

### Neural organoid networks on MEAs

Cerebral organoids were differentiated from human induced-pluripotent stem cells (hiPSCs) and cultured to generate diverse cell types within the tissue^2^. The identity of cells in the cerebral organoid was confirmed using immunohistochemistry. At day 115, we observed a mixed population of PAX6-positive progenitor cells, FOXG1-positive forebrain progenitor cells, TUBB3-positive neuronal cells and GFAP-positive astrocytes. In addition, we also observed the expression of more mature markers such as the deep layer marker CTIP2 as well as the later-born upper layer marker SATB2 (Figure S1A).

To analyze neuronal activity, the organoids were plated onto high-density multielectrode arrays (HD-MEAs). On a HD-MEA, one (solo) two (duo), or three (trio) organoids were plated. We cultured them for more than two weeks to grow their axons and form functional connections between the organoids (Figure 1A). Inter-organoid connections with axons were visually confirmed (Figure 1B). Within each organoid, synchronization activity was detected, suggesting that organoids form active and tightly connected networks inside them. Pairwise correlation analysis revealed that the synchronicity of the organoids was similar in solo, duo, and trio organoid networks (Figures 1C-E). Also, inter-organoid synchronization of their burst activity patterns was detected in duo and trio organoid networks (Figure 1C). Positive pairwise correlation between organoids revealed their synchronization. The inter-organoid correlation was weaker than the intra-organoid correlation in both duo and trio organoid networks, suggesting that the axonal connections between organoids were sparser than the intra-organoid synaptic connections (Figure 1D, E, Figure S1B and C). Together, these data suggests that our organoids are not only active but are also connected through axons, giving rise to a tissue network consisting of both local circuits within the organoids and inter-organoid connections across distal organoids.

**Figure 1.**
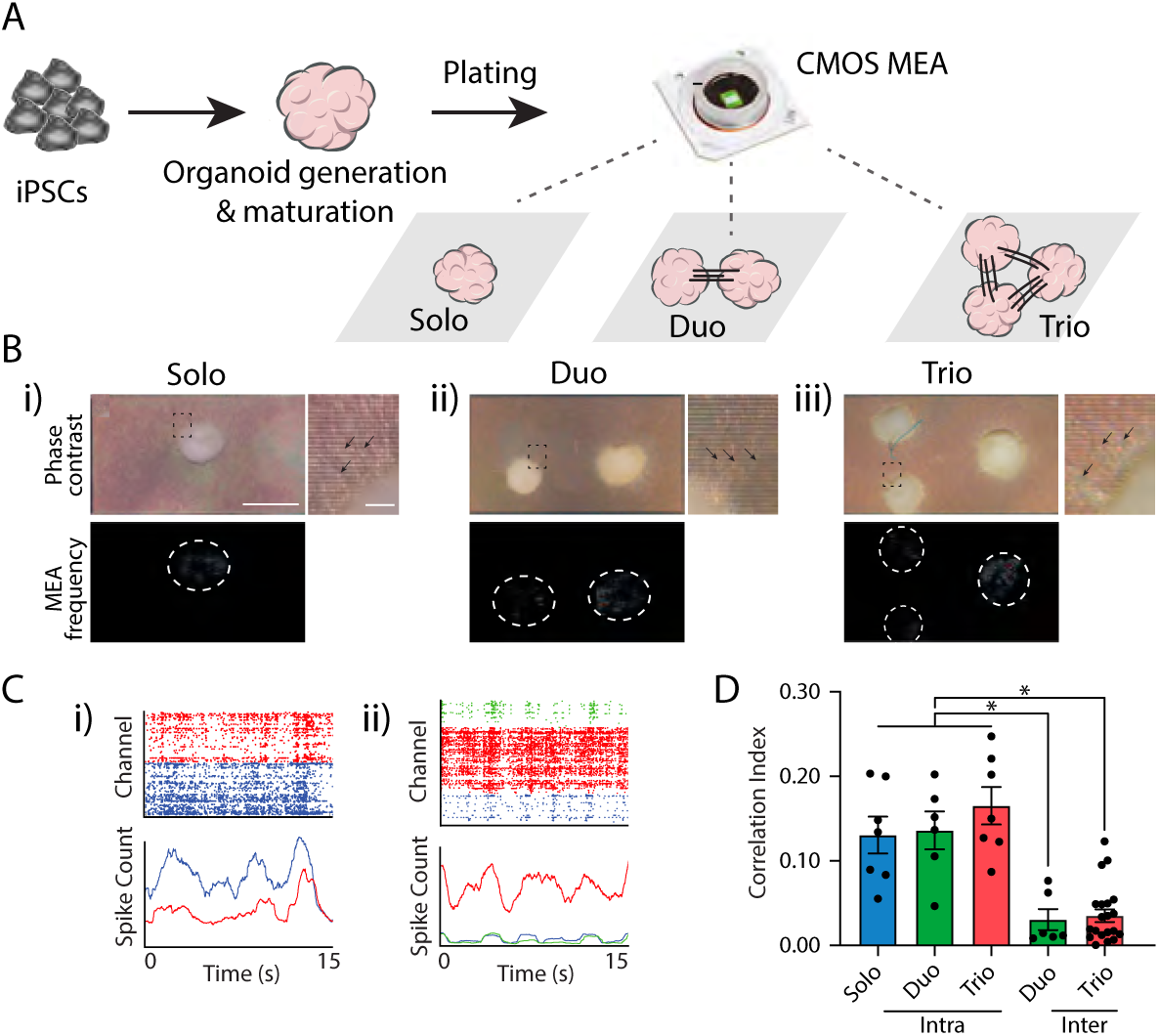
Characterization of solo, duo and trio organoid networks. (A) Schematic describing generation and plating of organoids. Organoids were generated from iPSCs and plated onto MaxOne HD-MEA chips to form solo, duo and trio organoids. (B) Images and activity scan of i) solo, ii) duo and iii) trio organoids on MaxOne chips. White dashed lines indicate active regions detected by activity scan. Inlet shows magnified images where arrows indicate axons. Scalebar = 1mm, 100µm (inlet). (C) Raster plot and correlation index of i) duo and ii) trio organoids showing synchronized activity between distal (interconnected) organoids. (D) Pairwise correlation index comparing synchronicity between solo, duo and trio organoids. Both intra-organoid and inter-organoid synchronicity was observed. *p < 0.05; one-way ANOVA followed by Tukey’s multiple-comparison test. Data are presented as mean ± SEM.

Stimulus decoding score increases in trio organoids following repetitive stimulation.

To address the functionality of organoids and their ability to respond to extrinsic signals, we set up a stimulus decoding task in which we stimulated at two distinct input locations and evaluated the output response at a different region within the network. With the help of computational algorithms, we asked whether it is possible to blindly discriminate between the two input stimulation patterns from the output signal (Figure 2A). Configurations of input and output regions were selected as following. In solo organoids, the two input locations and output region were chosen within the same organoid. For duo organoids, one organoid was assigned as the input organoid, and another as output organoid. The input organoid contained two stimulation electrodes apart from each other. For trio organoids, two organoids were assigned as the input organoids, while another organoid as output. Each input organoid contained one stimulation location (Figure 2A). The same stimulation parameter was applied to both stimulation 1 and 2, with a resting period between the two stimulations. Stimulation electrodes were chosen near active electrodes.

**Figure 2.**
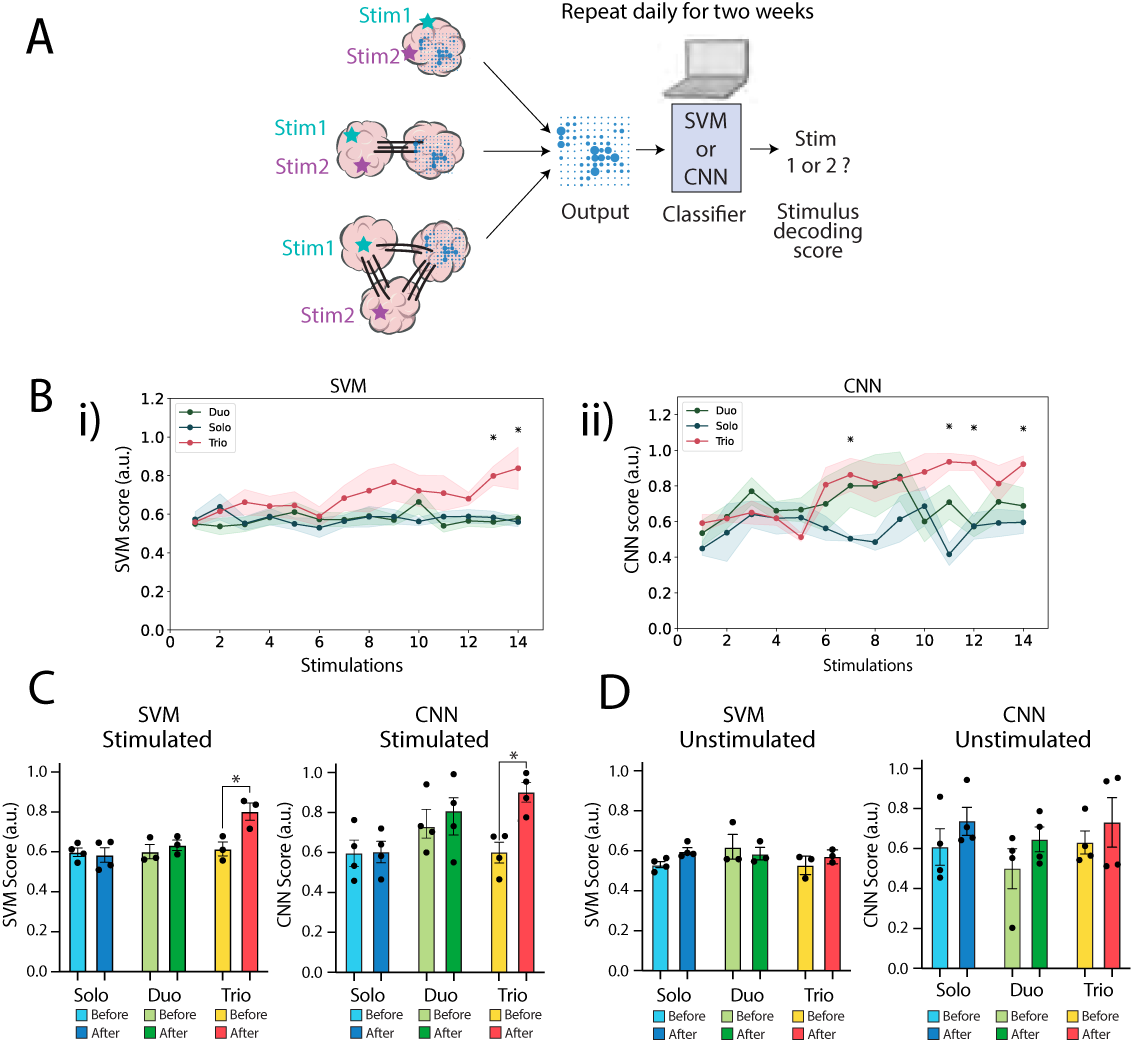
Stimulus decoding score increased in trio organoids following long-term stimulation. (A) Schematic showing setup of the stimulus decoding task. Input regions of solo, duo and trio organoids were stimulated daily for two weeks. The output response is fed into classification algorithms, and a stimulus decoding score (SVM/CNN score) is obtained. (B) Average i) SVM and ii) CNN score of solo, duo and trio organoids. Lines indicate mean ± SEM. *p < 0.05; one-way ANOVA followed by Tukey’s multiple-comparison test. (C) Average SVM and CNN score of solo, duo and trio organoids during the first half of stimulation (before) in comparison to the second half of stimulation (after). *p < 0.05; paired *t* test (two-sided). (D) SVM and CNN score at before and after two weeks of culture without daily stimulation. Data are presented as mean ± SEM.

Since we aimed to evaluate growth and changes of network activity by stimulus, we selected initial configurations in which the Support Vector Machine (SVM) algorithm is unable to distinguish between input stimulation 1 and 2. Initial rounds of stimulation failed to increase SVM discrimination score and did not induce any difference in evoked responses between stimulation 1 and 2 in the short-term. We thereby decided to extend our task to a daily stimulation over the period of two weeks to monitor long-term changes in discrimination score and neural activity.

Strikingly, we observed an increase in SVM score of trio organoids only following repeated, long-term stimulation (Figure 2B and S2A). Convoluted Neural Network (CNN) score also supports this results in which trio organoid consistently showed an increase in score over time (Figure 2B and S2B). Moreover, CNN results seem to suggest that duo organoids tend to perform better over solo organoids in some trials, implying that the bidirectional connection of duo organoids is also advantageous over solo organoids.

To understand whether long-term stimulation is necessary for the increase in stimulus decoding score, we assessed unstimulated organoids in which we did not stimulate the organoids daily as control. Unstimulated organoids were subjected to the stimulus decoding task once in week 0 and once again in week 2 to obtain initial and final stimulus decoding scores.

We confirmed that in early phases of stimulation (before training), solo, duo and trio organoids that were daily stimulated had a low stimulus decoding score. In later phases of stimulation (after training), trio organoids have a significantly higher average SVM and CNN score. In contrast, this increase was not observed in control, non-daily stimulated organoids, indicating that the increase in stimulus decoding score was not simply due to culture time, but a direct result of stimulation allowing the neurons to adapt and rewire (Figure 2E and F, Figure S2).

### Long-term network plasticity of organoids following long-term stimulation

The increase in stimulus decoding score in trio organoid networks following long-term stimulations suggests the importance of both intra- and inter-organoid connectivity in the functionality of organoids. To understand the fundamental mechanism underlying the surge in stimulus decoding score in trio organoids, we visualized and compared changes of evoked responses following stimulation 1 and 2 at the beginning of the stimulus decoding task (“before” training) and at the end of the two weeks (“after” training).

At the start of the regime, consistent with low stimulus decoding scores, we found that stimulation 1 and 2 elicited similar responses at output in trio organoids. After two weeks of stimulation, the spatiotemporal response patterns to stimulation 1 and 2 drastically differ, corresponding to the increased stimulus decoding score (Figure 3A-C). For example, evoked response maps showed two initial areas of output response following both stimulation 1 and 2. Although both areas remained responsive, long-term, repetitive stimulation reinforced a much stronger response from one area in response to stimulation 1 and another area to stimulation 2 (Figure 3A and B). Typically, stimulation responses peaked around 200ms following stimulation (Figure 3B). This increased separation following stimulation was not observed in solo and duo organoids. We were able to observe a decrease in standard deviation and propagation correlation of evoked response in trio organoids, indicating that both inter- and intra-organoid connectivity was significantly altered in trio organoids only (Figure 3C-E). These suggest that the stimulus-induced response of intra-organoid network within the output organoid is drastically modified by the long-term repetitive stimulation sessions.

**Figure 3.**
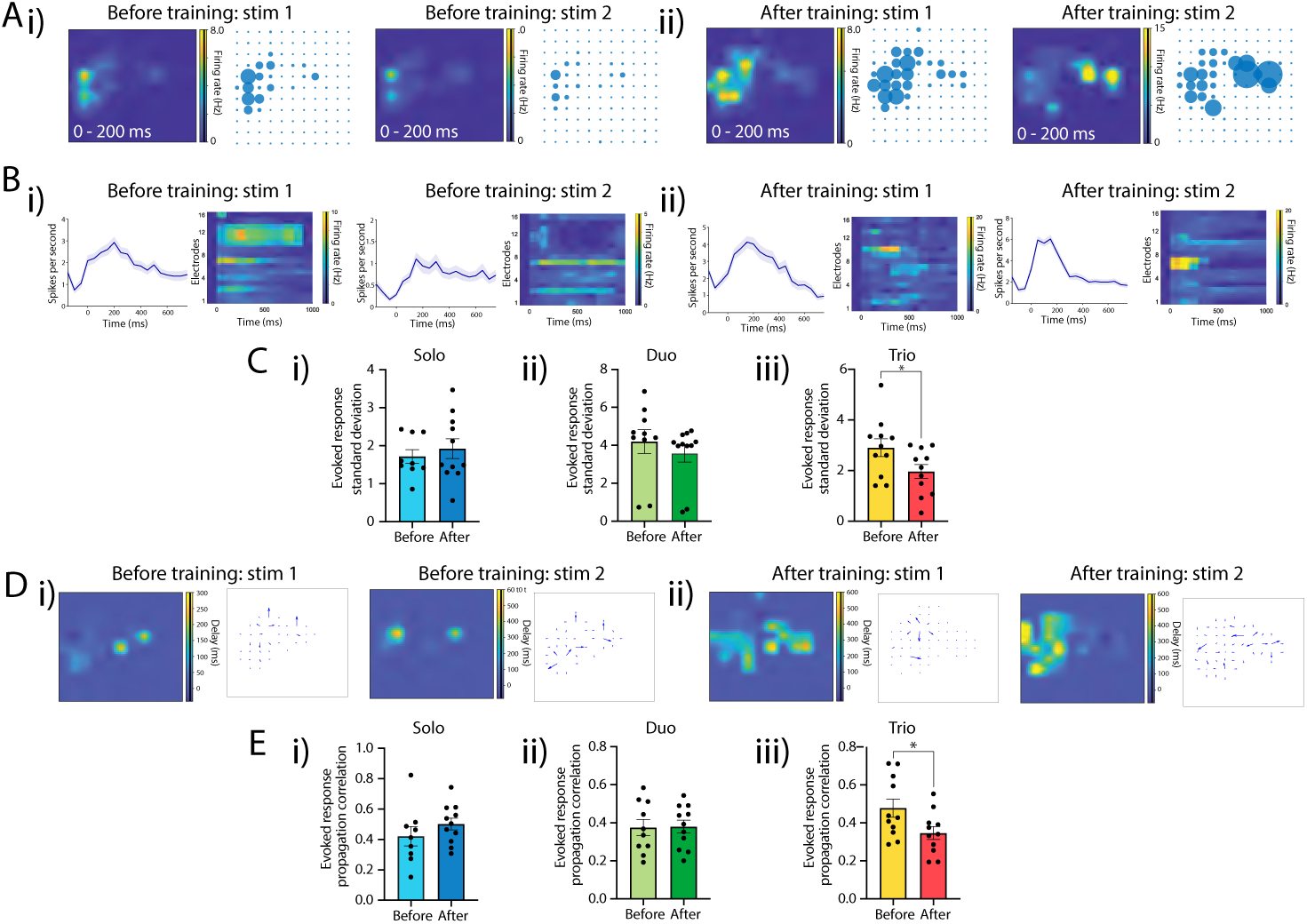
Intra-organoid response to stimulation changes following long-term stimulation. (A) Representative evoked activity maps showing response of output region 0-200ms following stimulation 1 and 2, i) during the beginning of the stimulus decoding task (before training) and ii) after two weeks of stimulus decoding task (after training). Repetitive stimulation led to differing response and increased segregation following stimulation 1 and 2. (B) Firing rate and heat map depicting responses following stimulation, i) before and ii) after training. Response curve (left) shows average spiking rate of all 121 output electrodes in response to stimulation over 800ms. Heat map (right) shows the top 16 most responsive electrodes following both stimulation 1 and 2. (C) Quantification of (B) showing standard deviation of all evoked response curves in i) solo, ii) duo and iii) trio organoids. Stimulation 1 and stimulation 2 were compared before and after training. A lower standard deviation suggests a less variable, more stable response. *p < 0.05; student t. test (two-way). (D) Heatmap of signal delay within output organoid, i) before and ii) after training. Vector plot (right) summarizes the direction and speed of delay. Arrows point towards direction of delay and arrow length denotes speed. Longer arrows indicate a faster speed and less delay. (E) Quantification of (D) showing correlation between the evoked response propagation of stimulation 1 and stimulation 2 in i) solo, ii) duo and iii) trio organoids before and after training. *p < 0.05; student t. test (two-way). Data are presented as mean ± SEM.

### Inter-organoid connectivity alteration following long-term stimulation

The network structure was further assessed by investigating alteration of spontaneous activity and propagation within the network (Figure 4). We detected neuronal activity that occurred in temporal proximity and calculated activity propagation directionality based on frequency and temporal delay. Notably, spontaneous propagation in trio organoid networks, but not solo or duo organoid networks, occurred more toward output organoid from the input organoids than the other direction after the long-term stimulation than before (Figure 4A). Correspondingly, activity propagation delay was reduced for the activity transmitted from input organoids to output organoids by the repeated stimulation sessions (Figure 4B). On the other hand, the network propagation delay on the other direction was increased (Figure 4C), suggesting that the network connectivity was biased towards the direction from input organoids to output organoids after the repetitive stimulations. Pairwise correlation analysis revealed that synchronicity between input and output organoid was increased after repetitive stimulations (Figure 4D), suggesting that the network connectivity between input and output organoids was induced by the stimulation sessions. Taken together, the repetitive stimulation induced connectivity from input to output organoids, implying that the input stimulation refined the inter-organoid connection and reshaped the network of trio organoids during the repeated stimulation sessions that increased the functionality.

**Figure 4.**
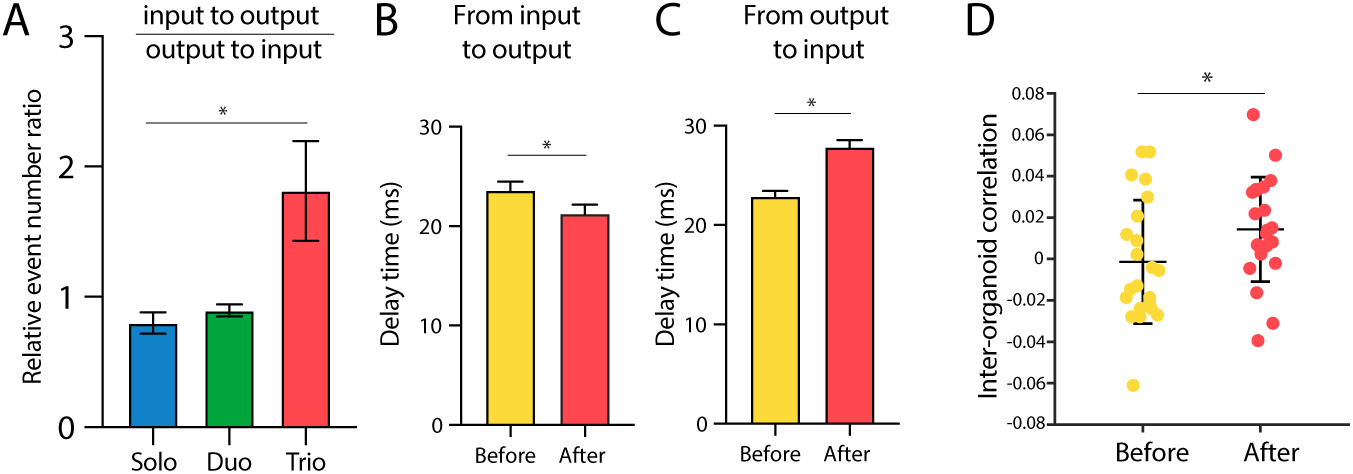
Spontaneous inter-organoid activity transmission is altered after long-term stimulation. (A) Ratio of number of events (spikes transmitted) from the output organoid to the input organoids relative to the number of events from the input organoids to the output organoid increased in trio networks after training normalized with the value before training. *p < 0.05; paired t. test (two-way). (B) Delay of transmission time is increased from output to input organoids in the trio network. *p < 0.05; paired t. test (two-way). (C) Delay of transmission time is decreased from input to output organoid in the trio network. *p < 0.05; paired t. test (two-way). Data are presented as mean ± SEM. (D) Pairwise correlation index comparing synchronicity between input and output organoid before and after stimulations. *p < 0.05; student t. test (two-way). Data are presented as mean ± STD.

## Discussion

In this study, we explored how connecting multiple neural organoids and subjecting them to repeated external stimulation can enhance their long-term functional refinement. Our major finding is that trio-organoid networks displayed substantially better stimulus-decoding capabilities over time than single or duo organoid networks, indicating that a more complex, hierarchical network structure can promote the development of robust input-output functionality. These results extend our current understanding of how in vitro neural systems can acquire functional attributes by receiving persistent and spatially distinct stimuli.

While many previous studies have focused on the immediate or short-term functional properties of cultured neurons^4,22,23^, we sought to emphasize the long-term mechanisms through which the networks of organoids develop the capacity to discriminate and encode signals. By monitoring and analyzing changes over two weeks of stimulation, we demonstrated how repetitive external stimuli can lead to the emergence of stable, decodable spatiotemporal patterns. This observation underscores that networks of organoids can gradually become specialized in how they process inputs—mimicking, to an extent, the developmental processes of the activity-dependent network refinement in the brain^9,24,25^.

Our findings also highlight the importance of hierarchical and modular networks in acquiring functionality. Connecting multiple organoids, each capable of forming its own local circuits, yielded a system in which distinct organoids could serve different roles, suggesting that hierarchical organization of the network is critical for the network functions as evident in the brain^26–28^. The modular architecture likely introduces variations in signal routing and plasticity, contributing to the enhanced decoding performance we observed. This is consistent with the previous studies demonstrated that modular arrangement of neuronal networks enhances functionality of the culture^29^.

Despite the promising outcomes of multi-organoid connectivity, several limitations remain. First, the total amount of stimulation in our experiments—100 times a day for two weeks—may still be insufficient to capture the full spectrum of plastic changes these organoids can undergo. Longer and more abundant stimulation could elicit additional adaptations or more stable circuit organization and should thus be tested in subsequent studies. Second, although our multi-organoid configuration yielded functional benefits, it is reported that a single organoid or cultured neurons on a two-dimensional surface could develop internal circuitry sufficient to perform certain stimulus-decoding tasks^4,22,30,31^. We suspect that our stimulation was not sufficient to trigger persistent alteration of the internal network for solo and duo organoid networks. Because we purposefully selected input-output electrode combinations that does not exhibit stimulus decoding ability with the relatively simple machine learning algorithms used in the study, it is possible that the experiment design negatively biased away from selecting circuit locations that can enhance network functionality with less and shorter stimulations. Further exploration of both single- and multi-organoid setups is warranted to isolate the conditions under which hierarchical networks become essential for diverse functional studies.

Testing the organoids with more complex signals and more sophisticated tasks will be critical. Presently, we used two clearly defined input locations to probe stimulus decoding. A next step is to increase the number or pattern of inputs (e.g., multi-frequency stimulation, spatiotemporal sequences) and assess how well organoid networks adapt to such complexity. Our assays were open-loop system which do not provide feedback to the organoids. As a next step, closed-loop set up could be also useful to evaluate the organoid network in the future. Additionally, we cannot yet pinpoint the exact cellular populations responsible for the long-term changes observed upon stimulation. Identifying potential “node cells” or hub neurons that connect within or between organoids—and conducting biochemical or molecular assays on those cells—could illuminate the mechanistic underpinnings of long-term synaptic plasticity and connectivity.

In conclusion, our work emphasizes that hierarchical network organization and role-specialization across multiple organoids can significantly enhance the emergence of long-term, stimulus-specific functionality in organoid-based neural circuits. By refining stimulation paradigms, extending culture times, and studying cellular adaptations at higher resolution, we can deepen our understanding of how to guide organoids toward brain-like information processing capabilities in vitro.

## Acknowledgement

This work was supported mainly by Research Institute of Advanced Technology, Softbank Corp., and in part by JSPS (20K20643, 24H02307, JPJSCCA 20190006); AMED-CREST (JP20gm1410001); HFSP (RGP012/2024) and the Institute for AI and Beyond. The study was also supported by JST SPRING (JPMJSP2108) and the ANRI fellowship.

## Declaration of Interests

The authors declare no competing interests

## Materials and methods

### Cell culture and organoid generation

409B2 human iPS cell line was obtained from Kyoto University through RIKEN Bioresource center cell bank (HPS0076)^32^. iPS cells were maintained in mTeSR Plus medium (STEMCELL Technologies) on Matrigel (Corning)-coated dishes. iPSCs were passaged at 70-80% confluency with ReLeSR (STEMCELL Technologies). All cells were cultured in a 5% CO2/37°C humidified incubator.

To generate cerebral organoids, iPSCs were dissociated into single cells using TrypLE Express (Thermo Fisher Scientific) and plated into 96 well-round bottom plates (Corning) at a density of 9,000 cells/well. Induction and generation of cerebral organoids were conducted using the STEMDIFF Cerebral Organoid Kit (STEMCELL Technologies). On day 7, the organoids were embedded in Matrigel Growth Factor-Reduced (Corning) and plated onto orbital shakers starting from day 10. Following a month of culture, the organoids were transferred into 10 cm dishes and the Maturation Medium was switched to a Maintenance Medium consisting of a 1:1 mixture of DMEM/F12 with Neurobasal medium supplemented with 2% B27 supplement with vitamin A, 1% N2 supplement, 1% GlutaMAX, 1% Penicillin-Streptomycin-Amphotericin (PSA) solution, 0.5% non-essential amino acids solution and 0.2% sodium bicarbonate. On day 100, the organoids were cut into smaller pieces using a scalpel and placed back into a 10 cm dish on an orbital shaker. The cut organoids were recovered for at least 10 days before plating onto MaxOne HD-MEA chips.

### Immunohistochemistry

Organoids were fixed in 4% PFA/4% sucrose solution overnight at 4°C. After two washes with PBS, the organoids were transferred into 30% sucrose and incubated overnight at 4°C. The organoids were embedded in OCT compound and subjected to cryosectioning. The tissues were incubated in Blocking Buffer (1xTBS containing 3% BSA, 5% NGS, 0.4% triton X-100 and 0.02% sodium azide) for 30 min at room temperature. Primary antibodies were diluted using the same solution and incubated overnight at 4°C. Samples were washed twice with PBS and incubated with secondary antibodies conjugated to Alexa Fluor 488 or 568 (Thermo Fisher Scientific) for one hour at room temperature.

The following primary antibodies were used: anti-CTIP2 (Abcam), anti-SOX2 (Cell Signaling Technologies), anti-TUBB3 (BioLegend), anti-FOXG1 (Abcam), anti-GFAP (Merck Millipore), anti-SATB2 (Abcam).

## Organoid networks on HD-MEA chips

Prior to use, MaxOne HD-MEA CMOS Chips (MaxWell Biosystems) were treated with 1% Tergazyme solution overnight. On the next day, the chips were rinsed 5 times with distilled water, sterilized in 75% ethanol for 10 minutes and fully dried before plating. Sterilized MaxOne Chips were coated with Matrigel Growth Factor Reduced diluted in DMEM/F12 (1:50) for one hour at room temperature.

One, two, or three organoids were placed onto the chip with minimal medium for an hour. Once the organoids attach and settle, 0.5 ml of Maintenance Medium was added to the plate. The plated organoids were cultured in a 5% CO2/37°C humidified incubator.

Over the next week, the medium was gradually changed to Recording Medium, consisting of BrainPhys Medium (STEMCELL Technologies) supplemented with 2% B27 Supplement with vitamin A, 1% N2 supplement, 1% GlutaMAX, 1% Penicillin-Streptomycin-Amphotericin (PSA) solution. Glucose concentration of the medium was adjusted to 20mM. Thereafter, a half medium change was done every 3-4 days.

## Determination of stimulation input and output

Electrical stimulation was performed through the MaxOne MEA system. For each plate, active regions of organoids were determined through the combination of activity scan results and organoid photos. A region of 121 electrodes (11 x 11, spacing of 5 electrodes between each electrode) was selected as the output region. Input stimulation electrodes were selected by test stimulating the most active electrodes determined by activity scan. Electrodes that can trigger a response in the output region were chosen as the two stimulation electrodes respectively.

### Repeated Stimulation

Repeated stimulation was applied to the organoid networks at two locations daily over the period of 2 weeks. One round of stimulation consists of 100 times of bi-phasic stimulation delivered every second at one location. Within each stimulation, ten of 800 mV, 200 μs pulses were applied at 100 Hz. The organoid networks were rested for a hundred seconds between two sessions of stimulation for two locations.

### Data analysis

The recorded data was analyzed using MATLAB. Signals were bandpass filtered with a frequency range of 300-3000 Hz, whereas peaks with a value more than 5 times the standard deviation was identified as a spike. Correlation index within and between organoids was calculated using Pearson correlation coefficient function corrcoef(). For the calculation within an organoid pairwise correlation of spike activity (binned at 10 ms) between all electrodes was performed, whereas for inter organoid calculation pairwise correlation of summed spike activity (binned at 10 ms) between each organoid was used. For evoked responses, spike activity in the 900 ms time window after each stimulation for each electrode was extracted for stimulation 1 and stimulation 2. For evoked response map, mean number of evoked spikes in the first 200ms was extracted for each electrode and plotted. For network response, histogram of spike timings for evoked spikes from all stimulations were plotted with a 50 ms binning. Activity delay map was calculated using finddelay() function, by pairwise comparison of spike activity between each electrodes. Propagation direction was derived from the delay map using gradient() function and plotted using quiver() function.

### Stimulus decoding by classification score

Neural activity data recorded for 500 ms immediately following each electrical stimulation was collected using the MaxLab Python API to train a classifier model for each trial. The classifier models were created and trained separately for each trial, while their hyperparameters were fixed and shared across all trials. The classification accuracy of the models was evaluated to determine how distinguishable the neural activities became in the corresponding trial in response to the two stimulation patterns. Two Python-based supervised machine learning algorithms were employed to classify the post-stimulation signals: a support vector machine (SVM) and a deep learning convolutional neural network (CNN).

The Support Vector Machine (SVM) classifier was implemented using the Scikit-learn library with a radial basis function (RBF) kernel. The data was compressed into a one-dimensional format before being input to the SVM classifier. The accuracy of the classifiers was evaluated using Scikit-learn’s default scoring function.

The Convolutional Neural Network (CNN) classifier was established using the PyTorch-based Braindecode library^33^. The model employed a learning rate of 0.0625*0.01, the default 2D CNN-based ShallowFBCSPNet layer architecture, a negative log-likelihood loss function (NLLLoss), the Adam optimizer with weight decay, and a cosine annealing strategy for accuracy evaluation. For the CNN classifier, the two-dimensional data, consisting of channel information and raw electrical signal across time, was directly fed into the model.

## Notes

### Competing Interest Statement

The authors have declared no competing interest.

## References

1 Eiraku, M. et al. Self-organized formation of polarized cortical tissues from ESCs and its active manipulation by extrinsic signals. Cell Stem Cell 3, 519–532 (2008). 10.1016/j.stem.2008.09.002

2 Lancaster, M. A. et al. Cerebral organoids model human brain development and microcephaly. Nature 501, 373–379 (2013). 10.1038/nature12517

3 Kagan, B. J. et al. Toward a nomenclature consensus for diverse intelligent systems: Call for collaboration. Innovation (Camb) 5, 100658 (2024). 10.1016/j.xinn.2024.100658

4 Cai, H. et al. Brain organoid reservoir computing for artificial intelligence. Nature Electronics 6, 1032–1039 (2023). 10.1038/s41928-023-01069-w

5 Morales Pantoja, I. E., et al. First Organoid Intelligence (OI) workshop to form an OI community. Front Artif Intell 6, 1116870 (2023). 10.3389/frai.2023.1116870

6 Tau, G. Z. & Peterson, B. S. Normal development of brain circuits. Neuropsychopharmacology 35, 147–168 (2010). 10.1038/npp.2009.115

7 Stiles, J. & Jernigan, T. L. The basics of brain development. Neuropsychol Rev 20, 327–348 (2010). 10.1007/s11065-010-9148-4

8 Katz, L. C. & Shatz, C. J. Synaptic activity and the construction of cortical circuits. Science 274, 1133–1138 (1996). 10.1126/science.274.5290.1133

9 Pan, Y. & Monje, M. Activity Shapes Neural Circuit Form and Function: A Historical Perspective. J Neurosci 40, 944–954 (2020). 10.1523/JNEUROSCI.0740-19.2019

10 Hübener, M. & Bonhoeffer, T. Neuronal plasticity: beyond the critical period. Cell 159, 727–737 (2014). 10.1016/j.cell.2014.10.035

11 Jamann, N., Jordan, M. & Engelhardt, M. Activity-dependent axonal plasticity in sensory systems. Neuroscience 368, 268–282 (2018). 10.1016/j.neuroscience.2017.07.035

12 Mathis, M. W., Perez Rotondo, A., Chang, E. F., Tolias, A. S. & Mathis, A. Decoding the brain: From neural representations to mechanistic models. Cell 187, 5814–5832 (2024). 10.1016/j.cell.2024.08.051

13 Velasco, S. et al. Individual brain organoids reproducibly form cell diversity of the human cerebral cortex. Nature 570, 523–527 (2019). 10.1038/s41586-019-1289-x

14 Sharf, T. et al. Functional neuronal circuitry and oscillatory dynamics in human brain organoids. Nature Communications 13, 4403 (2022). 10.1038/s41467-022-32115-4

15 Samarasinghe, R. A. et al. Identification of neural oscillations and epileptiform changes in human brain organoids. Nat Neurosci 24, 1488–1500 (2021). 10.1038/s41593-021-00906-5

16 Chow, S. Y. A., Hu, H., Osaki, T., Levi, T. & Ikeuchi, Y. Advances in construction and modeling of functional neural circuits in vitro. Neurochem Res 47, 2529–2544 (2022). 10.1007/s11064-022-03682-1

17 Meunier, D., Lambiotte, R. & Bullmore, E. T. Modular and hierarchically modular organization of brain networks. Front Neurosci 4, 200 (2010). 10.3389/fnins.2010.00200

18 Sporns, O. & Betzel, R. F. Modular Brain Networks. Annu Rev Psychol 67, 613–640 (2016). 10.1146/annurev-psych-122414-033634

19 Onesto, M. M., Kim, J. I. & Pasca, S. P. Assembloid models of cell-cell interaction to study tissue and disease biology. Cell Stem Cell 31, 1563–1573 (2024). 10.1016/j.stem.2024.09.017

20 Osaki, T. et al. Complex activity and short-term plasticity of human cerebral organoids reciprocally connected with axons. Nat Commun 15, 2945 (2024). 10.1038/s41467-024-46787-7

21 Zhang, Z., O’Laughlin, R., Song, H. & Ming, G. L. Patterning of brain organoids derived from human pluripotent stem cells. Curr Opin Neurobiol 74, 102536 (2022). 10.1016/j.conb.2022.102536

22 Kagan, B. J. et al. In vitro neurons learn and exhibit sentience when embodied in a simulated game-world. bioRxiv (2021).

23 Yada, Y., Yasuda, S. & Takahashi, H. Physical reservoir computing with FORCE learning in a living neuronal culture. Applied Physics Letters 119, 173701–173701 (2021).

24 Agetsuma, M. et al. Activity-dependent organization of prefrontal hub-networks for associative learning and signal transformation. Nat Commun 14, 5996 (2023). 10.1038/s41467-023-41547-5

25 Bragg-Gonzalo, L., De León Reyes, N. S. & Nieto, M. Genetic and activity dependent-mechanisms wiring the cortex: Two sides of the same coin. Semin Cell Dev Biol 118, 24–34 (2021). 10.1016/j.semcdb.2021.05.011

26 Zamora-López, G., Chen, Y., Deco, G., Kringelbach, M. L. & Zhou, C. Functional complexity emerging from anatomical constraints in the brain: the significance of network modularity and rich-clubs. Sci Rep 6, 38424 (2016). 10.1038/srep38424

27 Müller-Linow, M., Hilgetag, C. C. & Hütt, M. T. Organization of excitable dynamics in hierarchical biological networks. PLoS Comput Biol 4, e1000190 (2008). 10.1371/journal.pcbi.1000190

28 Betzel, R. F. & Bassett, D. S. Multi-scale brain networks. Neuroimage 160, 73–83 (2017). 10.1016/j.neuroimage.2016.11.006

29 Yamamoto, H. et al. Impact of modular organization on dynamical richness in cortical networks. Sci Adv 4, eaau4914 (2018). 10.1126/sciadv.aau4914

30 Isomura, T., Kotani, K., Jimbo, Y. & Friston, K. J. Experimental validation of the free-energy principle with in vitro neural networks. Nat Commun 14, 4547 (2023). 10.1038/s41467-023-40141-z

31 Akita, D., Suwa, E., Ikeda, N. & Takahashi, H. Neural Activity and Information Processing Capacity of Neuronal Culture. Annu Int Conf IEEE Eng Med Biol Soc 2023, 1–4 (2023). 10.1109/EMBC40787.2023.10340168

32 Okita, K. et al. A more efficient method to generate integration-free human iPS cells. Nat Methods 8, 409–412 (2011). 10.1038/nmeth.1591

33 Gramfort, A. et al. MEG and EEG data analysis with MNE-Python. Front Neurosci 7, 267 (2013). 10.3389/fnins.2013.00267

