## Supplementary figures for "Repetitive stimulation modifies network characteristics of neural organoid circuits"

Siu Yu A. Chow *et al.*

**This PDF file includes:**

Figs. S1 to S3

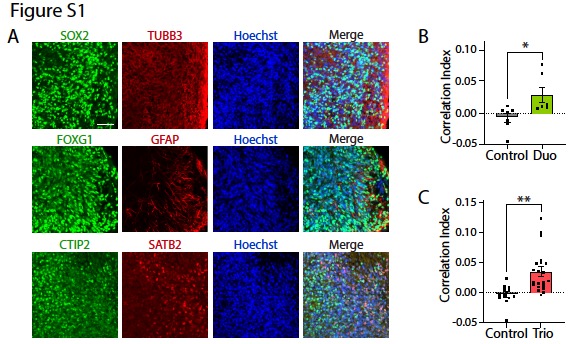

**Fig S1. Characterization and activity of solo, duo and trio organoids**

(A) Immunostaining of organoids at day 115. Scalebar = 50 μm.
(B) Pairwise correlation index comparing duo organoids on the same chip with random, unrelated organoids. *p < 0.05; student’s *t* test (two-sided).
(C) Pairwise correlation index comparing trio organoids on the same chip with random, unrelated organoids. *p < 0.05; student’s *t* test (two-sided).

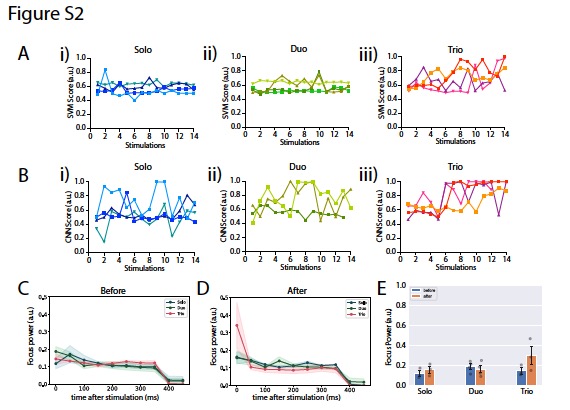

**Fig. S2. Focus power of CNN in solo, duo and trio organoids**

(A) Stimulus decoding SVM score of i) solo, ii) duo and iii) trio organoids.
(B) Stimulus decoding CNN score of i) solo, ii) duo and iii) trio organoids.
(C) Focus power of solo, duo and trio organoids after stimulation over time, where total focus power of 0-500ms = 1.0. Each datapoint represents a binning of 50ms as a percentage of the total focus power.
(D) Focus power of solo, duo and trio organoids after stimulation over time.
(E) Quantification of (A) and (B) comparing the focus power between solo, duo and trio organoids before and after stimulation (0-50ms post stimulation).

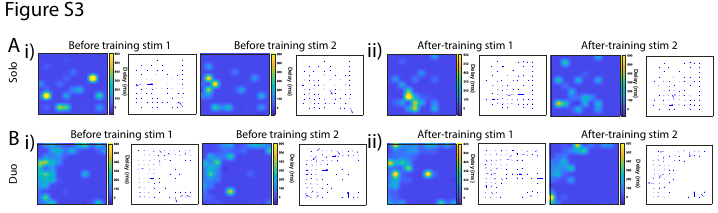

**Fig. S3. Output signal delay of solo and duo organoids**

(A) Heatmap of output signal delay (solo organoid).
(B) Heatmap of output signal delay (duo organoid).
